# Life-long persistence of infectious Zika virus: inflammation and behavioral sequela

**DOI:** 10.1101/2020.06.11.145854

**Authors:** Derek D.C. Ireland, Mohanraj Manangeeswaran, Aaron Lewkowicz, Kaliroi Engel, Sarah M. Clark, Adelle Laniyan, Jacob Sykes, Ha Na Lee, Ian McWilliams, Logan Kelly-Baker, Leonardo H. Tonelli, Daniela Verthelyi

**Affiliations:** US Food and Drug Administration, Office of Biotechnology Products; University of Maryland School of Medicine, Department of Psychiatry

## Abstract

The recent spread of Zika virus (ZIKV) and its association with congenital defects and neurological disorders has created an urgent need to understand the pathogenesis of ZIKV and identify therapeutic strategies that will prevent or eliminate them. The neurodevelopmental defects associated with ZikV infections early in pregnancy are well documented, however the potential defects associated with infections in late pregnancy and perinatal period are less well characterized. Further, the long-term sequelae of these infections are not fully understood. Immunocompetent C57BL/6 mice infected at one day old (P1), which neurodevelopmentally model late pregnancy in humans, develop a transient neurological syndrome including unsteady gait, kinetic tremors, severe ataxia and seizures 10-15 days post-infection (dpi) but symptoms subside after a week, and most animals survive. Despite apparent recovery, MRI of convalescent mice shows reduced cerebellar volume that correlates with altered coordination and motor function as well as hyperactivity and impulsivity. Persistent mRNA levels of pro-inflammatory genes including *Cd80, Il-1α*, and *Ifn-γ* together with *Cd3, Cd8* and perforin (*PrfA)*, suggested the persistence of a low-grade inflammatory process. Further, the brain parenchyma of convalescent mice harbor multiple small foci with viral antigen, active apoptotic processes in neurons, and cellular infiltrates, surrounded by activated astrocytes and microglia as late as 1-year post-infection. Detection of negative-sense strand viral RNA and replication of virus derived from these convalescent mice by blinded passage in Vero cells confirmed that low levels of live ZikV persist in CNS of convalescent mice life-long. Our studies establish that Zika can establish reservoirs in CNS and suggest that anti-viral treatment that clears virus from the CNS as well as long-term neurological and behavioral monitoring may be needed for patients known to be exposed to ZikV at an early age.

**Author’s summary:** The congenital brain malformations associated with ZikV infections early in pregnancy are well documented, however whether apparently asymptomatic perinatal exposure could lead to long term sequelae is not fully understood. Using a non-lethal neonatal mouse model, we examine host-pathogen interactions, anatomical changes and behavioral patterns by following survivors of the acute infection for over 1 year. We discover that infectious Zika virus has the potential to remain in the CNS for life, lodged within small foci surrounded by gliosis and infiltrating immune cells that may act to limit the viral spread, but also interfere with healing and contribute to life-long neuropathic and behavioral sequelae. These results suggest that anti-viral treatment and long-term neurological and behavioral monitoring may be indicated for patients known to have been exposed to Zika virus, regardless of neurodevelopmental disease severity.

## Introduction

Zika virus (ZIKV) is an arbovirus from the Flaviviridae family that caused a world-wide public health emergency between 2014 and 2016 as over 750,000 cases were reported in 84 countries, including over 40,000 in the US(1). Although transmission of ZIKV has since declined, WHO now considers Zika endemic to many areas of the world and infection clusters continue to be reported in regions of Cuba, India and Southeast Asia, where there are large populations of women of childbearing age who are susceptible to the virus (2-4),(5). Zika infections are asymptomatic in most adults, but approximately 20 % of subjects develop-fever, rash, conjunctivitis, muscle and joint pain (6). In a small number of cases ZikV infections have been linked to more serious neurological disease including: Guillain Barre Syndrome, acute myelitis, encephalitis and polyneuropathies, as well as chorioretinitis(7, 8). Importantly, infections are more detrimental during early pregnancy as ∼10 % of children born to mothers infected during the 1^st^ trimester show developmental defects at birth including microencephaly, lissencephaly, cerebral atrophy, ventriculomegaly, cerebellar hypoplasia, brain calcifications, microphthalmia, and arthrogryposis (9), (10),(11), (12). These defects have been collectively termed Zika congenital syndrome. Among the children of women exposed during the second and third trimester, the incidence of Zika congenital syndrome was lower, but there are multiple reports of neurodevelopmental defects, hydrocephaly, seizures, hearing, motor, behavioral and sociability deficits, suggesting that Zika’s pathological effect extends beyond the early developmental period (13). Importantly these defects can be evident at birth or develop later in life suggesting that the full neurological and behavioral impact of this disease may become apparent as these children grow(13, 14). Further, reports of virus in the spinal fluid months after birth suggest a delay in full viral clearance from the CNS (15). Thus, it is critical to gain a better understanding of the host-pathogen interactions, both during acute infection and after apparent recovery, to identify possible interventions to reduce the developmental impact and minimize the damage that viral persistence could cause.

To better understand the long-term effects of ZIKV infections our group developed a model using immunocompetent C57BL/6 mice (B6) challenged at one day after birth (P1) with ZikV (PRVABC059 strain, subcutaneously) (16). Previous studies comparing human and mouse CNS developmental markers such as stages of neurogenesis, axon extension, establishment and refinement of synapsis, and myelin formation, indicate that a P1 mouse brain corresponds to the human brain during the second to third trimester stage of development (17, 18). We have previously shown that P1, C57BL/6 mice infected with ZikV develop a transient encephalitis characterized by unsteady gait, kinetic tremors, severe ataxia, loss of balance and seizures that manifest 10-12 days post infection (dpi) and subside by 26-30 dpi (16). CNS infection is accompanied by inflammation, cellular infiltration predominantly of CD8+ T cells, and increased mRNA expression levels of interferon-gamma (*Ifn-γ*), granzyme B (*GzmB*), and perforin (Prf1), that is associated with neurodegeneration and loss of Purkinje cells (16). Over 80% of the mice survive the challenge and live a normal life span, without displaying ataxia, seizure or other overt signs of neurological deficit. The non-lethal outcome allows for the study of the natural progression of ZIKV infection in a developing brain and any resulting structural and functional deficits. Importantly, this mouse model bypasses transplacental transmission and consequent placental insufficiency, which can also impact on neurological development.

Recent studies assessing the effects of Zika infections showed that mice can have behavioral and motor deficits as late as 60 dpi (19). Moreover, in a recent study we showed that the chorioretinitis that affects these mice is evident as late as 90 dpi (20). In this study, we follow the natural history of the surviving mice s and show that although the signs of acute encephalitis subside, the mice have significant life-long deficits in motor function, coordination and anxiety that correlate with reduced brain volume, particularly affecting the cerebellum. We further show that live, replicating Zika virus persists in the brain of surviving mice for more than 1 year, and is associated with ongoing apoptosis, within foci that are surrounded by activated astrocytes and microglia. The relative contribution of the original tissue damage and the persistent ZikV replication and surrounding inflammation in the CNS, to the observed neurological deficits in adults remains unclear, however it suggests that anti-viral drugs may be indicated to completely eradicate the virus from the brain and that anti-inflammatory or immunosuppressive therapies in patients that were infected with Zika early in life should be undertaken with caution. Further, the data indicated the need to closely monitor exposed children, as an apparently transient ZikV infection early in development may result in life-long behavioural and psychological deficits. Lastly, our studies provide a platform to test therapeutics to clear ZikV from the CNS and/or to mitigate damage caused by the associated chronic inflammation.

## Results

### Acute Zika infection in CNS in neonatal mice

Subcutaneous infection with ZikV in neonatal (P1) C57BL/6 mice leads to a transient viremia, followed by infection of the CNS (Figure 1A)(16, 20). As previously published, the virus is cleared from peripheral blood and most organs within 9 dpi, but in the CNS viral RNA peaks at 6-12 dpi and persists long after symptoms have resolved (Figure 1A) (16) (21). At 12 dpi, IF-IHC shows ZIKV antigen primarily in the cortex, hippocampus, and cerebellum (Figure 1B), however animals recover and by 30 dpi little residual viral antigen can be detected, even in tissues that had been heavily infected such as the cerebellum (Figure 1B). The clinical evolution of the encephalitis follows the viral load and the symptoms, which include reduced weight gain, ataxia, dystonia, and seizures, resolve by 24 dpi (16) (22) (23) (24). Overall, fewer than 20 % of the animals succumb to infection and surviving animals have a normal lifespan with higher than 80% surviving by 12 months. (Figure 1A).

**Figure 1:**
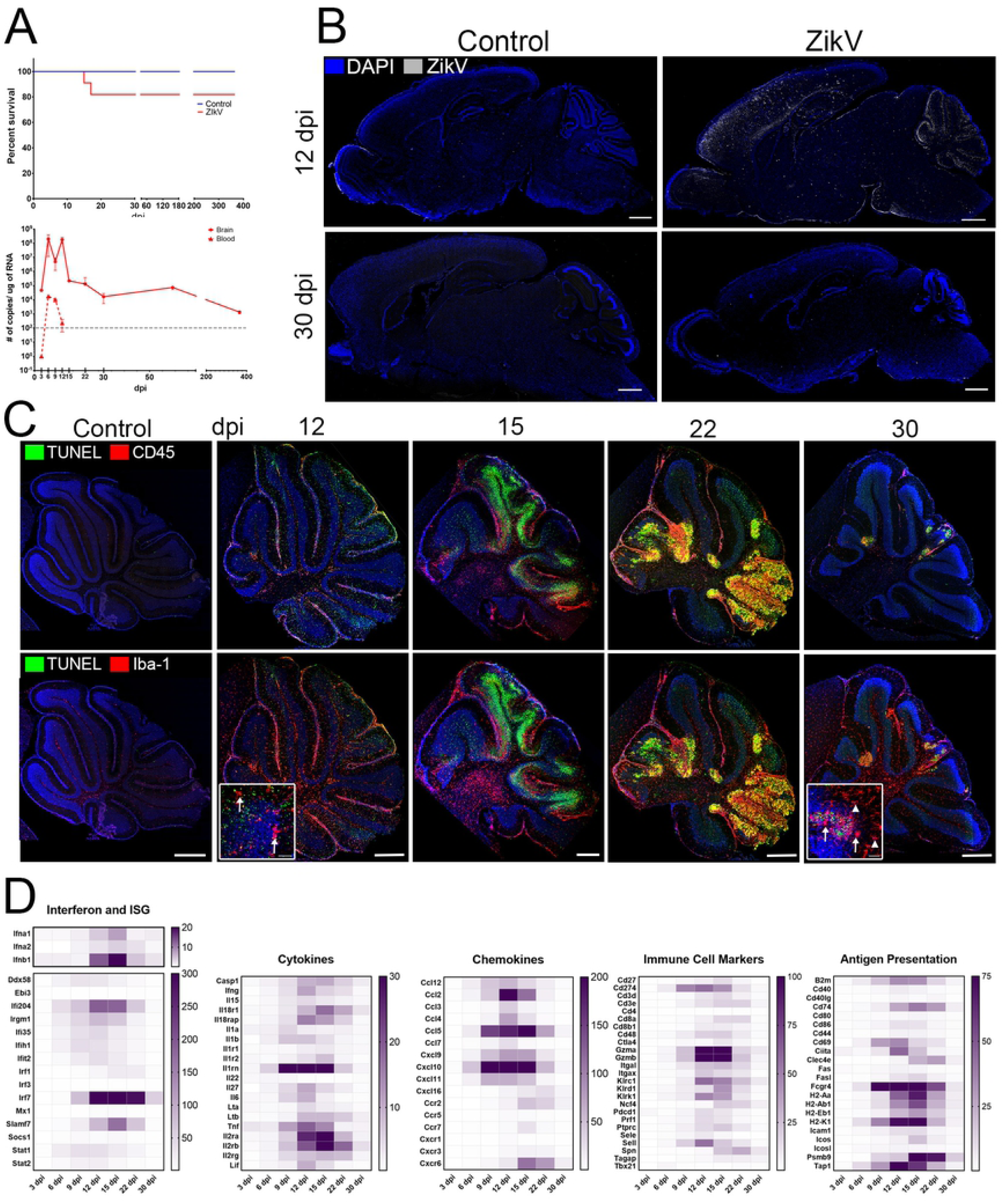
Acute ZikV infection in CNS. Neonatal C57Bl/6 mice were challenged with 2000 TCID_50_ SC on P1. **A**. Top. Survival curve. 20 % of animals succumb to disease by 18 dpi, 80% survive (n > 20/group). Bottom. Viral RNA copies/μg RNA in CNS and blood. By 15 dpi (peak of disease) ZikV RNA levels are significantly reduced relative to peak levels (9 – 12 dpi) (n = 6 mice/timepoint). After reduction, viral RNA is detectable in the CNS throughout life of the animal. **B**. Sagittal brain sections of ZikV infected and aged-match controls stained for ZikV. Image shows representative image of 4-6 animals per time point; scale bar = 1 mm. **C**. Cerebellum of ZikV infected animals at indicated dpi. Top. Staining for CD45+ infiltrating immune cells (red) and apoptosis (TUNEL, green). Bottom. Staining with TUNEL (green) and Iba-1+ microglia (red) indicating microgliosis. Both infiltrating immune cells and activated microglia progressively colocalize with distinct apoptotic foci in the cerebellum. Scale bar = 500 μm. Insets. Confocal maximum projection of TUNEL and Iba-1+ microglia in granular layer of cerebellum. Arrows indicate activated microglia. Arrow heads indicated resting microglia, distal from apoptotic foci at 30 dpi. Scale bar = 50 μm. **D**. Heatmap of RNA expression (fold change, n = > 6 infected, n = > 3 uninfected from at least 2 independent experiments per time point in the brain of ZikV-infected animals, from 3 dpi to 30 dpi, as detected by Nanostring technology. Genes selected based on significant changes during disease course, from a panel of 596 genes and organized by gene function. A full table is available upon demand.

During the acute infection there is significant meningitis and swelling of the pia as well as extensive cellular infiltration with increased numbers of CD45^+^ lymphocytes and Iba-1^+^ activated microglia evident in the infected areas, particularly in cerebellum, (Figure 1C and Supplementary Figure 1). Assessment of immune-related gene expression suggests that inflammation peaks at 15 dpi, as there is increased expression of genes linked to interferon responses (*Ifnα, Ifnβ, Irf7, Ifi204, Slamf7*); recruitment and activation of macrophages and neutrophils (*Ccl2, Ccl5, Cxcl10, Cxcl11, Cxcr2, Cxcr6)*; antigen presentation (MHC (*H2-Aa, -Ab1, - Eb1, -K1*), *B2m, Cd74, Tap1, Psamb9, Cd274, Cd80, Cd86*), and inflammation (*C3, Tnf, Il1, Il18, Il6, and Casp1*), as well as genes that suggest the presence of infiltrating T and NK cells (*Cd3, Cd4 and Cd8, Il2, Gzma/b and Prf1, Klrc1, Kirk1, and Ifnγ*) (Figure 1D). The infected areas show significant TUNEL staining indicating extensive apoptosis and neurodegeneration in cerebellum that results in a disorganization of cerebellar cortex and reduced number of Purkinje cells (Figure 1 C and Supplementary Figure 2A), although areas of apoptosis are also evident in the occipital cerebral cortex and hippocampus (supplementary Figure 2B). Importantly, as the viral load decreases and the symptoms subside, cellular infiltration and gliosis in cerebellum resolve, and inflammation of the pia diminishes (Figure 1C and Supplementary Figure 1). Together, these data suggested that ZikV causes a symptomatic but transient meningoencephalitis in neonatal mice.

### Increased levels of fluid in CNS and reduced cerebellum volume in convalescent mice

While ZikV-infected mice older than 30 days showed no obvious tremors, ataxia or seizures, the extensive tissue disruption during acute disease suggested that the transient infection could result in permanent loss of cerebral mass or calcifications paralleling the lesions described for exposed human offspring. To evaluate whether the infection led to permanent structural damage, we performed magnetic resonance imaging (MRI) of infected mice at 60 dpi (“young”), 4-6 (“adult”), and 12-14 (“aged”) months post-infection (mpi). As shown in Figure 2, at 60 dpi the overall volume of the brain was smaller compared to controls, consistent with recently published findings (19). Of note, as the mice age and catch-up in body weight (Supplementary Figure 3), the differences in total brain volume relative to age-matched controls became less evident suggesting that the tissue repairs over time (Figure 2B). However, convalescent animals showed increased percentage fluid (as indicated by white) in the CNS, especially in and around cerebellum and a reduction in cerebellar volume that was evident at all the time points. This fluid increase appears to be associated with expansion of the 4^th^ ventricle and/or the thickness of the pia layers surrounding the cerebellum but may also reflect changes to the microstructural composition of the tissue. Therefore, although animals that survived neonatal ZikV infection with PRVABC059 do not exhibit overt microcephaly, they do have reduced cerebellar volume and increased cerebellar fluid. Importantly, multiple studies have shown that defects in cerebellum can affect not only balance and coordination, but language, emotion, social behavior and working memory as well (25, 26).

**Figure 2:**
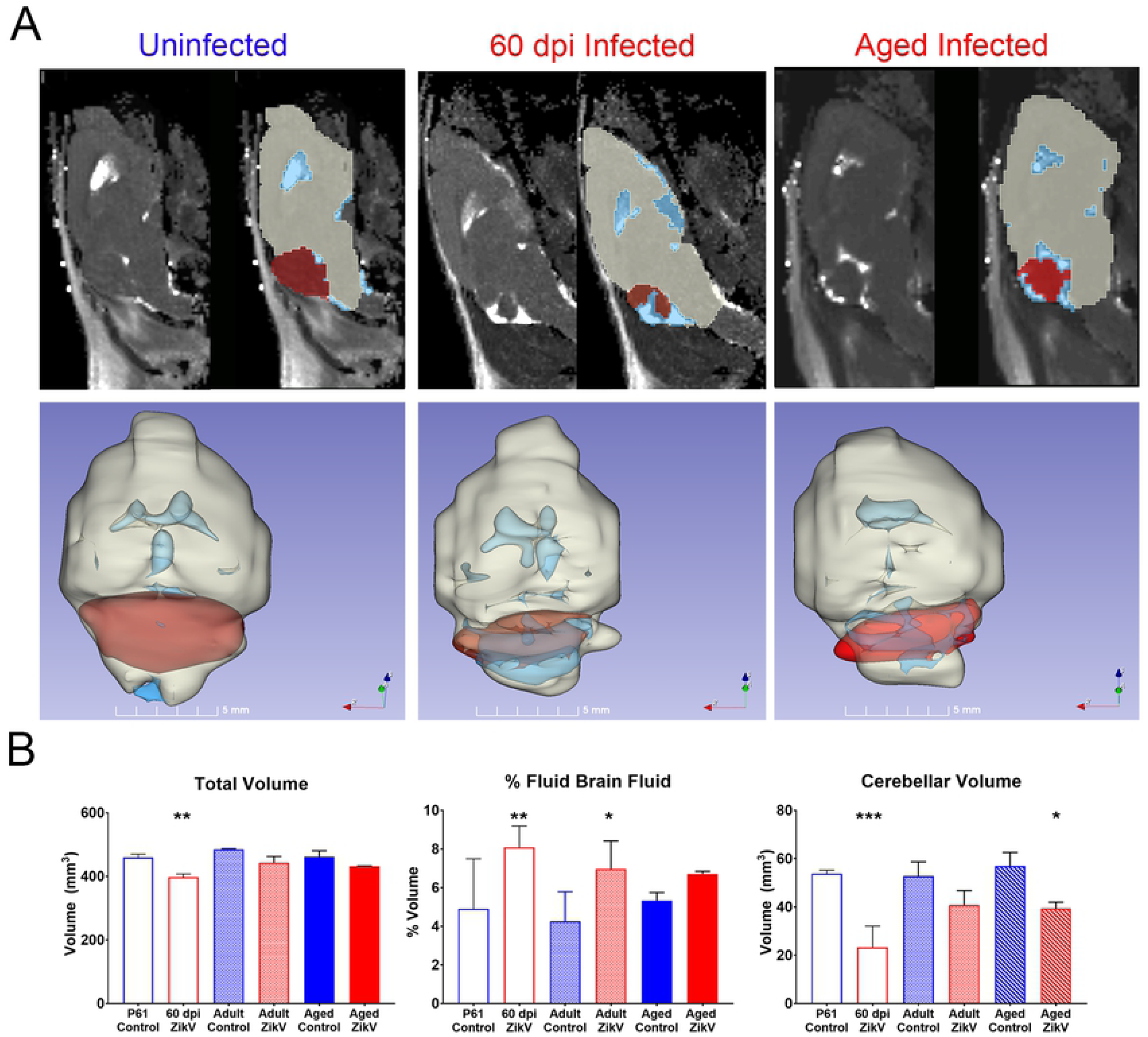
Magnetic Resonance Imaging of ZikV infected brains. **A**. representative examples of 2D slices of T2 MRI scans performed on uninfected, 60 dpi and > 1 ypi ZikV-infected animals. Representative 3D reconstructions of 2D slices, constructed using 3D Slicer software. The cerebellum is in red and fluid in the CNS in blue. **B**. Quantification of MRI for brain volume, % fluid in brain and cerebellum for each age group in the study, as analysed using the 3D Slicer software (N = 6 control, >7 infected mice per time point).

### ZikV infection results in life-long motor and behavior defects

Despite the reduction in cerebellar volume, the behavior of convalescent mice in their home cage environment was grossly indistinguishable from age-matched control mice. To examine more closely whether the early infection resulted in functional sequelae, we tested locomotor activity, as well as social, emotional and learning behavior of convalescent mice in three age groups: “young adult” (60 dpi), “adults” (4-6 months post infection (mpi)) and “aged” (> 1-year post infection (ypi)) mice. All animals were exposed to the same battery of 5 tests, under the same conditions and in the same environment, by a blinded researcher. The rotarod test, which examines motor function and coordination (Figure 3A) showed a significant reduction in latency to fall among ZikV-infected mice for all ages compared to age-matched controls, revealing an impairment in motor coordination and endurance (U = 253; *p < 0*.*0001;* Figure 3).

**Figure 3:**
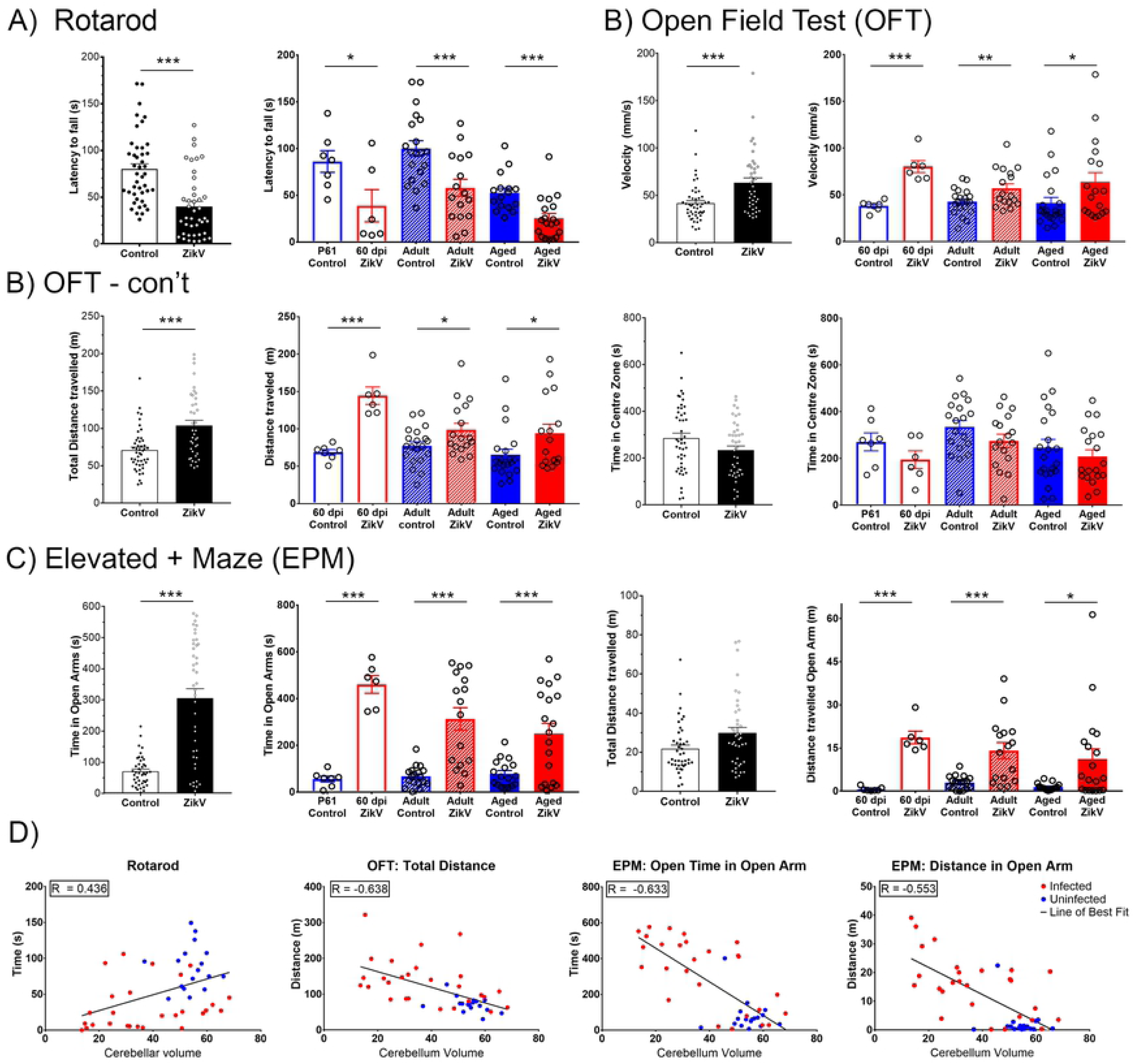
Behavioural studies in convalescent mice. For each test data from infected vs. uninfected mice are shown in black and white bars, while the red and blue bars show the same data parsed by age at the time of testing: young adults (60 dpi); adult (4-8 mpi) and aged (12-14mpi). **A**. Rotarod assay shows time (latency to fall) balancing on a rotating rod of gradually increasing speed over 180 seconds. **B**. Open Field Test (OFT) shows velocity, total distance travelled in the OFT maze and time within the center 20 cm^2^ of the maze over a 30-minute period. **C**. Elevated Plus Maze (EPM), explores time and movement in the different areas of the maze. Shown are time spent and distance travelled in the open arm. **D. Correlations of performance in behavioral studies and MRI measurements**. See Supplementary Table 1 for a full list of correlations. Statistical analysis for behavioral studies were conducted using both R (R Core Team (2019) and GraphPad Prism (v. 8.0.1: San Diego, CA). Spearman’s rank correlation coefficient was used to correlate MRI and behavior scores. (N = 48 controls, N = 42 ZikV).

Assessment of motility and exploratory behavior using the Open Field Test (OFT) showed that total horizontal motor activity and velocity were increased in ZikV-infected mice regardless of age, suggesting a hyperactive phenotype (U = 409, *p = 0*.*005*; Figure 3B). The increased mobility was more pronounced in younger mice (p<0.005), but evident at all stages (p<0.05). Interestingly, while convalescent mice covered more distance in the OFT, they did not spend more time than the control mice in the center of the arena, suggesting that the emotional component of this test was not affected (Figure 3B, Supplementary Figure 3B). However, ZikV-infected mice spent significantly more time (p<0.001) and traveled longer distances (p<0.001) on the open arm of the Elevated Plus Maze (EPM) than on the closed arm when compared to age-matched controls regardless of age (U = 247; *p < 0*.*0001*) (Figure 3C, Supplementary Figure 3B). Indeed, a significant proportion of ZikV infected mice, across all ages, spent greater than 50% of their time in the open arm, suggesting that ZikV-convalescent mice have reduced anxiety and thigmotactic tendencies. The time spent in the open arm did not correlate with the total distance travelled, indicating that the amount of time in the open arms does not reflect an overall increase in total locomotor activity, but a change in behavioral risk processing. Remarkably, other measures of sociability and learning such as the Social Interaction tests and Novel Object Recognition (Supplementary Figure 3C, D) did not show significant differences between convalescent mice and controls.

Having observed that ZikV convalescent mice had shorter latency times to fall from the rotarod, displayed hyperactivity in the OFT, and spent more time in the open arms of the EPM (p ≤ 0.001), we next assessed whether the magnitude of the behavioral changes correlated with the structural changes evident in MRI (Figure 2). As shown in Figure 3E, significant correlations were evident between the brain fluid content and cerebellum volume and latency to fall, as well as between cerebellar volume and the distance travelled in the OFT time or the time and distance traveled in the open arm of the EPM (see full list of correlations in Supplementary Table 1). Together, these data indicated that the reduced cerebellar size and increased fluid surrounding the cerebellum of infected animals persists throughout the life of the animals and correlate with motor and behavioral changes.

### ZikV induced neural damage persists long after acute disease

In this infection model, during the acute infection, the virus localizes primarily to glutamic acid decarboxylase-positive (GAD67^+^) neurons in the hippocampus, occipital cortex, and most prominently cerebellar cortex, and the infection results in increased apoptosis and neurodegeneration, particularly of Purkinje cells and other neurons in the granular layer (16) (Figure 1B, Supplementary, Figure 4, and Supplementary Figure 2A). Since Purkinje cells are critical for the regulation and coordination of motor movement, we next determined whether the infection caused permanent damage to the cellular structure of the cerebellum. Staining of neurons, including GABAergic Purkinje cells, with Neurofilament (NF-H, cocktail of mAb clones SMI-31 and SMI-32) and GAD67 in convalescent mice showed a persistent degradation of the Purkinje cell layer (PCL), both at 60 dpi and 1ypi, throughout the cerebellum (Figure 4A). In addition, the staining shows an increase in dystrophic and damaged neurites as well as a reduction in the density and complexity of neurites, especially in the molecular layer of the cerebellum of convalescent mice (Figure 4B). This indicated that the cerebellar pathology induced by ZikV is permanent and could contribute to the motor and coordination defects described above.

**Figure 4:**
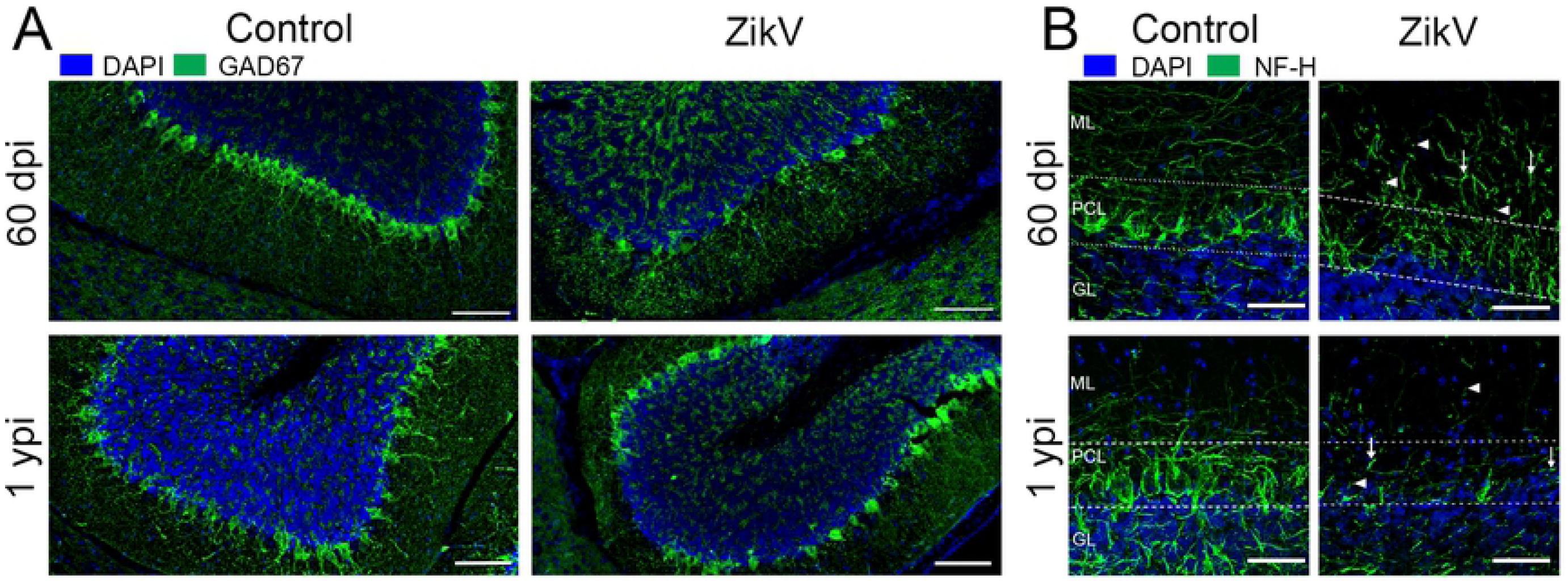
Neuropathology in the cerebellum following neonatal ZikV infection. **A**. GAD-67 staining (green) for GABAergic neurons in cerebellum of infected and uninfected controls. Scale Bars = 100 μm. **B**. Confocal imaging of neurofilament heavy chain (NF-H, green) staining in cerebellum. Images are maximum intensity projections of at least 16 Z-planes through each section. ML = molecular layer, PCL = Purkinje cell layer, GL = Granular Layer. White arrows indicate dystrophic neurites, white arrowheads indicate interrupted neurites. Scale Bar = 50 μm.

### Increased cellular infiltration in convalescent mice

As shown above, 30dpi mice had significant reduction in viral load that was associated with fewer infiltrating cells and reduced apoptosis, as well as lower levels of most of the immune-related genes that had been upregulated during the acute infection. This suggested that the infection and associated inflammation was resolving. Despite this, small foci of apoptosis and immune cells in and around the mid-brain, hippocampus and in the granular and molecular layers of the cerebellum were still present at 60 dpi. These foci concentrated in regions that were heavily infected during the acute disease (Figure 1, supplementary Figure 5,).

Confocal imaging of the cerebellum, showing TUNEL+ NeuN+ cells indicated that at least some of the cells undergoing apoptosis within these foci were neurons (Figure 5 and Supplementary Figure 6A). Further, CD45^+^ immune cells were found in proximity to and within these foci, which were surrounded by GFAP^+^ astrocytes and rounded Iba-1^+^ activated microglia (Figure 5 and supplementary Figure 6A). Surprisingly, similar apparently active foci of infection could be detected throughout the life of the mice as shown for the 1-year old mice, although the older mice seem to have fewer NeuN+ cells in the center of the foci. Of note, while the microglia immediately surrounding the foci have hypertrophic bodies and asymmetric processes, indicating activation, those that are distant to these apoptotic foci are in a resting state, with small soma and long, complex processes, suggesting that the areas of inflammation are very localized (Figure 5 and Supplementary Figure 6B). Together this showed the life-long presence of localized, active inflammatory processes in the brains of convalescent mice.

**Figure 5:**
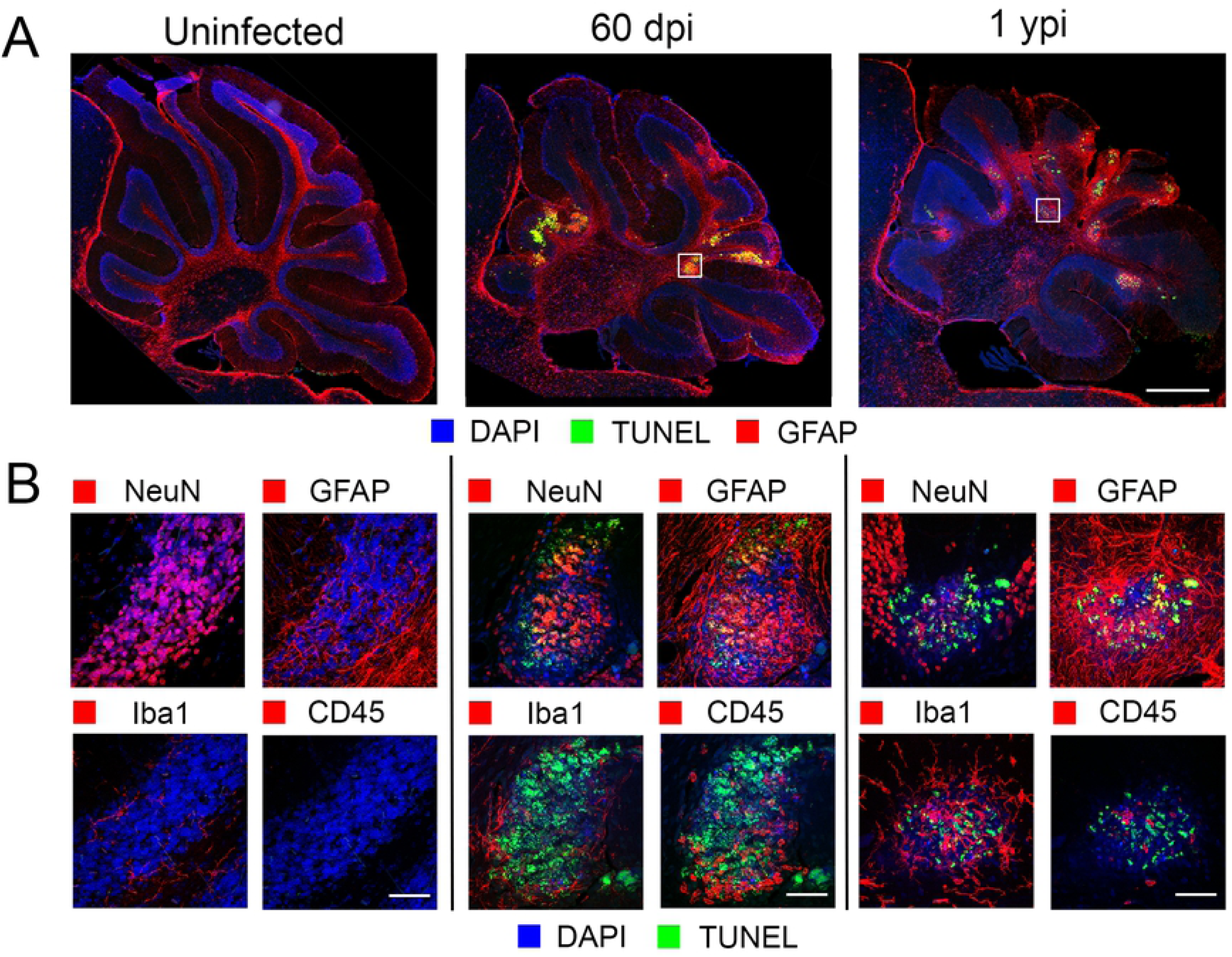
Apoptotic foci in cerebellum of ZikV infected animals. **A**. IHC staining of cerebellum of infected mice 60 dpi and 1ypi shows apoptotic foci (TUNEL^+^, green; DAPI, blue) surrounded by activated astrocytes (GFAP^+^, red). Scale Bar = 500 μm. **B**. Confocal microscopy of these lesions indicates apoptotic cells (TUNEL+, green and DAPI, blue in all images), surrounded by activated astrocytes (GFAP+, red) with activated microglia (Iba-1, red) and infiltrating immune cells (CD45+, red) proximal to and within the lesions. Scale Bars = 50 μm.

### Persistent inflammatory gene expression in CNS of ZikV-exposed animals

Acute meningoencephalitis due to ZIKV was associated with the up-regulation of the expression of genes linked to interferon responses, recruitment and activation of macrophages, and neutrophils, inflammation, as well as genes corresponding to infiltrating T cells (Figure 1D). To investigate whether the small foci of infiltrating cells and activated microglia were associated with an increase in the expression of immune-related genes, we examined the mRNA expression level for a panel of immune-related genes in cerebellum and cerebrum of convalescent mice at 2, 4-6 and 12-14 mpi (Figure 6). Surprisingly, while the relative expression levels were lower than during acute infection, the brains of convalescent animals showed a similar pattern of gene expression with increased levels of interferon-stimulated genes such as *Cxcl10, Cxcl11*, and *Oas1*. The expression of genes linked to macrophage/microglial activation including: *H2-Eb1, Cd40, Cd68, Cd80*, and *Cd86*, as well as *C3, Il6* and *Tnf* were also increased, regardless of age. In addition, a marked increase in chemokine expression, particularly *Ccl*2 and *Ccl5*, as well as pro-inflammatory cytokines *Ifnγ* and *Il1α* was evident at all ages. Lastly, in accordance with the increased CD45^+^ cells shown by IHC (Figure 5, Supplementary Figures 5, 6), there were increased levels of *Cd3, Cd8a, Ctla4, Gzmb, Icos* and *Gitr*, suggesting the presence of residual or bystander CD8+ T cells in the tissue and confirming the presence of active inflammatory processes in the CNS months after the challenge. Interestingly, the increased levels of mRNA were detectable not only in cerebellum, where the foci are located, but also in cerebral cortex suggesting that many areas of the parenchyma are exposed to increased inflammatory signals long after initial infection.

**Figure 6:**
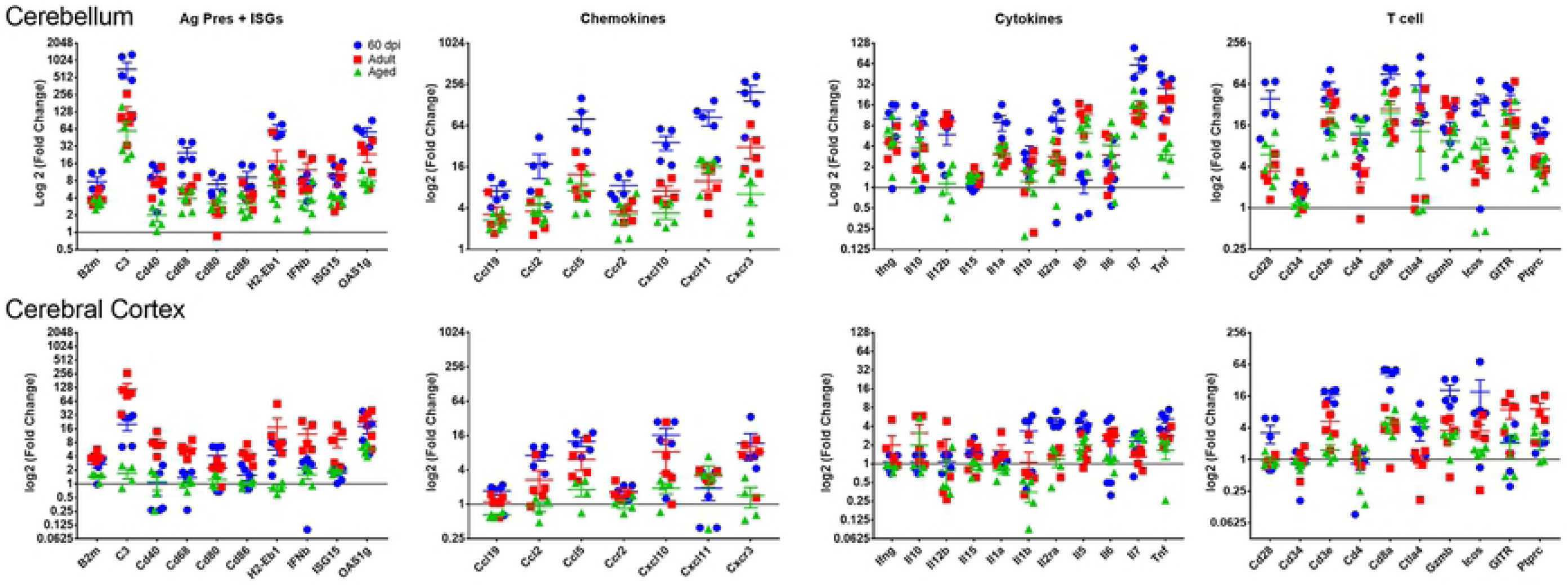
Expression of immune-related genes in the CNS of ZikV infected animals. Cerebellum and cerebral cortex were isolated from the CNS of 60-day (blue symbols), adult (4-8 mo., red symbols) and aged (> 1y, green symbols) ZikV-infected animals and gene expression analysed by real-time PCR using Mouse Inflammation Taqman array cards (Thermo Scientific). Genes associated with antigen presentation (Ag Pres) and IFN-inducible genes (ISG); T cell function; chemokines and pro-inflammatory cytokines; were grouped together. Ct values analysed using Quant-studio software (Thermo Scientific) using relative threshold algorithm for analysis. Data is presented as normalized against house-keeping genes and fold-change over age-matched, mock-infected controls as calculated with the ΔΔCt method.

Testing gene expression in the subgroup of mice that had undergone the MRI and behavioral testing allowed us to correlate the expression of pro-inflammatory genes with the behavioral sequelae in individual mice. The distance covered in OFT as well as the time and distance covered in the open arm of the EPM correlated with the mRNA levels in cerebellum for pro-inflammatory genes including *Cd80, Cd86*, MHC, and *Cd68*, ISGs C*xcl10, Cxcl11*, and *Stat1*, and pro-inflammatory markers *C3, Csf1, Il-1α, Il-18*. In addition, cerebellum size correlated with the expression levels of *Ifnγ, Tbet, Stat1, Stat4, Prf1 and Gzmb* suggesting an association with Th1 linked responses, while changes to the ratio of social to non-social interaction correlated with levels of *Il-14 (r>0*.*7)* and *Il-13 (r= 0*.*48)* in cortex. In addition, reduced latency to fall in the rotarod correlated with increased expression of *Tbet, Icos* and *Cd4*, although not *Cd8* (Supplementary Table 2). Chronic inflammatory processes in CNS have been associated with Parkinson’s, Alzheimer’s, and other neurodegenerative diseases of aging. Future studies will need to explore the extent to which the protracted inflammatory process contributes to the motor and behavioral deficiencies and whether it could predispose convalescent subjects to neurodegenerative diseases later in life.

### ZikV RNA, ZIKV antigen, and infectious ZIKV persists in the CNS of neonatally infected animals

Initial experiments using TCID_50_ assay on homogenates from the hemispheres of infected animals were negative after 22 dpi suggesting that infectious ZikV had been cleared from the CNS. Further, staining with human polyclonal antibody or mouse monoclonal antibody specific for the E-protein for ZikV antigen showed vestigial viral antigen in the cerebellum by 30 dpi (Figure 1) and no positive staining in cerebellum or other regions of the brains of exposed mice by 60 dpi (not shown). However, the persistent inflammatory gene expression, including type 1 IFN and ISGs, as well as the ongoing apoptosis, infiltration of immune cells and gliosis occurring at foci within the cerebellum of adult animals suggested that low levels of virus, below the level of detection of two-step IHC, could persist in the CNS of neonatally infected animals. To address this, we amplified the IF-IHC signalling using a multi-step technique with hyper-peroxidase labelled, secondary antibodies combined with a fluorescent HRP substrate, to improve the detection of residual ZikV antigen. As shown in Figure 7A, viral antigen was evident within foci of the granular layer (as marked by activated microglia) of the cerebellum at both 60 dpi and 1 ypi suggesting that the inflammatory processes surrounded the residual viral antigen.

**Figure 7:**
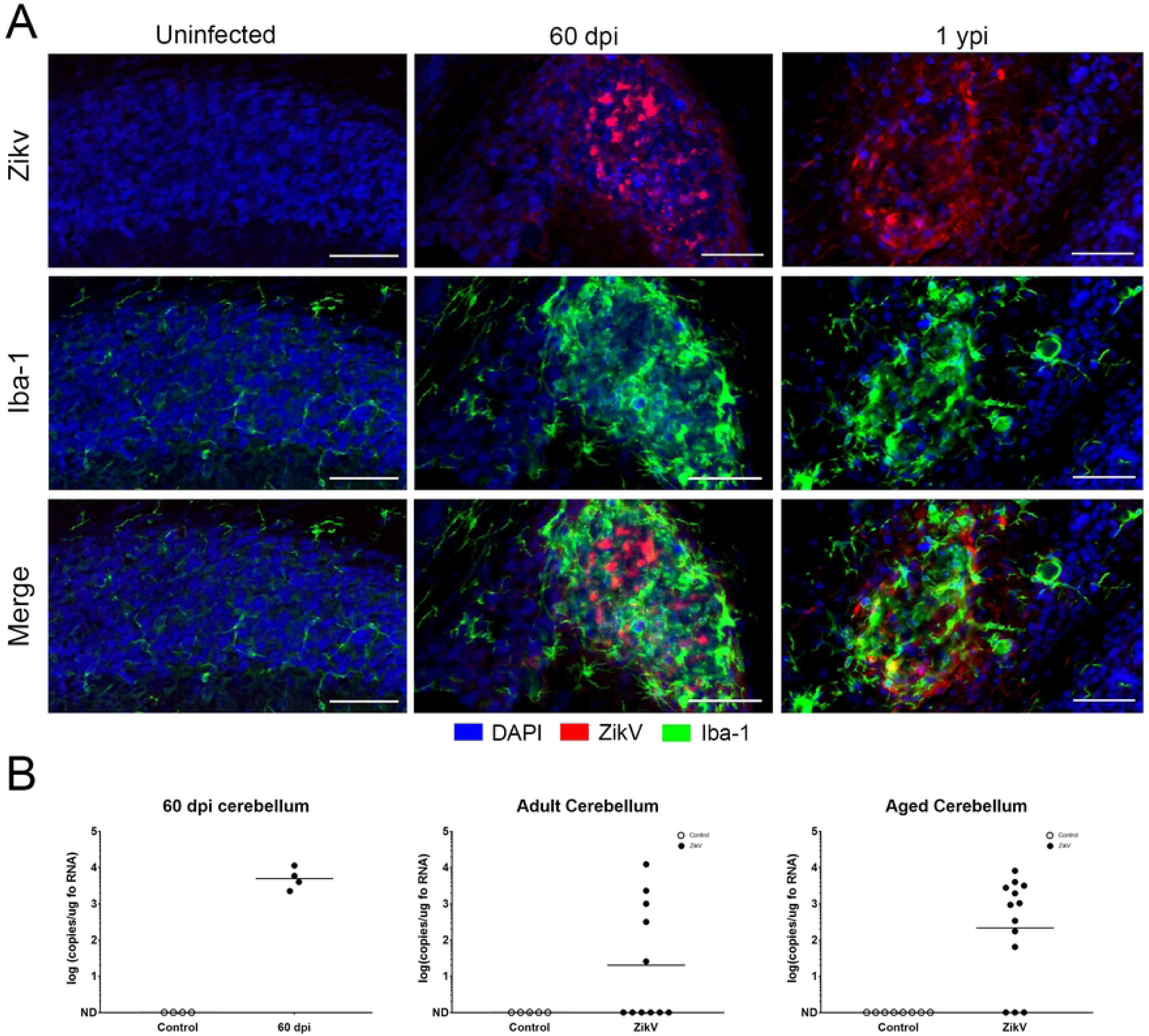
ZikV antigen and RNA persistence within cerebellar lesions. **A**. IF-IHC using Tyramide Signal amplification in combination with anti-ZikV E protein (red, top) identifies virus antigen within cerebellar lesion at 60 dpi and 1 ypi. Note the hypertrophied nuclei of activated Iba+ microglia (green) surrounding the viral antigen (bottom, merge). Scale Bars = 50 um. **B**. Real-time PCR for viral genome in cerebellum at 60 dpi, 4-8 mpi and 12-14mpi.

To determine whether the viral antigen was associated with viral RNA, we next tested the tissue for viral RNA using a one-step RT-qPCR, with primer/probes specific to detect ZikV. This showed the presence of viral genome RNA (positive-sense) in the brains of mice at 60 days 4 to 6 months, and 1 year post infection in 40-60% of convalescent mice (Figure 7B and Supplementary Fig 7). ZikV expression levels appeared higher in cerebellum (10^3^ – 10^5^ copies of viral RNA/ug total RNA) than cortex (Supplementary Figure 7, 10^1^ – 10^3^ copies of viral RNA/ug total RNA). To confirm that the RNA detected was not residual but represented replicating virus, we next tested the brains that had positive-sense ZikV RNA for the presence of negative-sense RNA strands. Negative-sense RNA is considered indicative of ongoing replication as it is a replicative intermediate of positive-sense RNA viruses and typically has a very short-half life. All the mice with quantifiable levels of viral RNA (positive-sense strand) showed detectable levels of negative-sense strand RNA, confirming the presence of replicating virus (Figure 8A). Lastly, we examined whether the detectable virus in convalescent adults was infectious by seeding brain homogenates on VERO cells, which after 5-7 days were fixed and stained with anti-ZIKV E-protein. As shown in Figure 8B, VERO E6 cells exposed to the cerebellar homogenates of mice older than 1 ypi, served to replicate ZIKV indicating infection of these cells by virus. This confirmed that ZIKV can establish a life-long reservoir in the CNS of mice challenged early in development.

**Figure 8:**
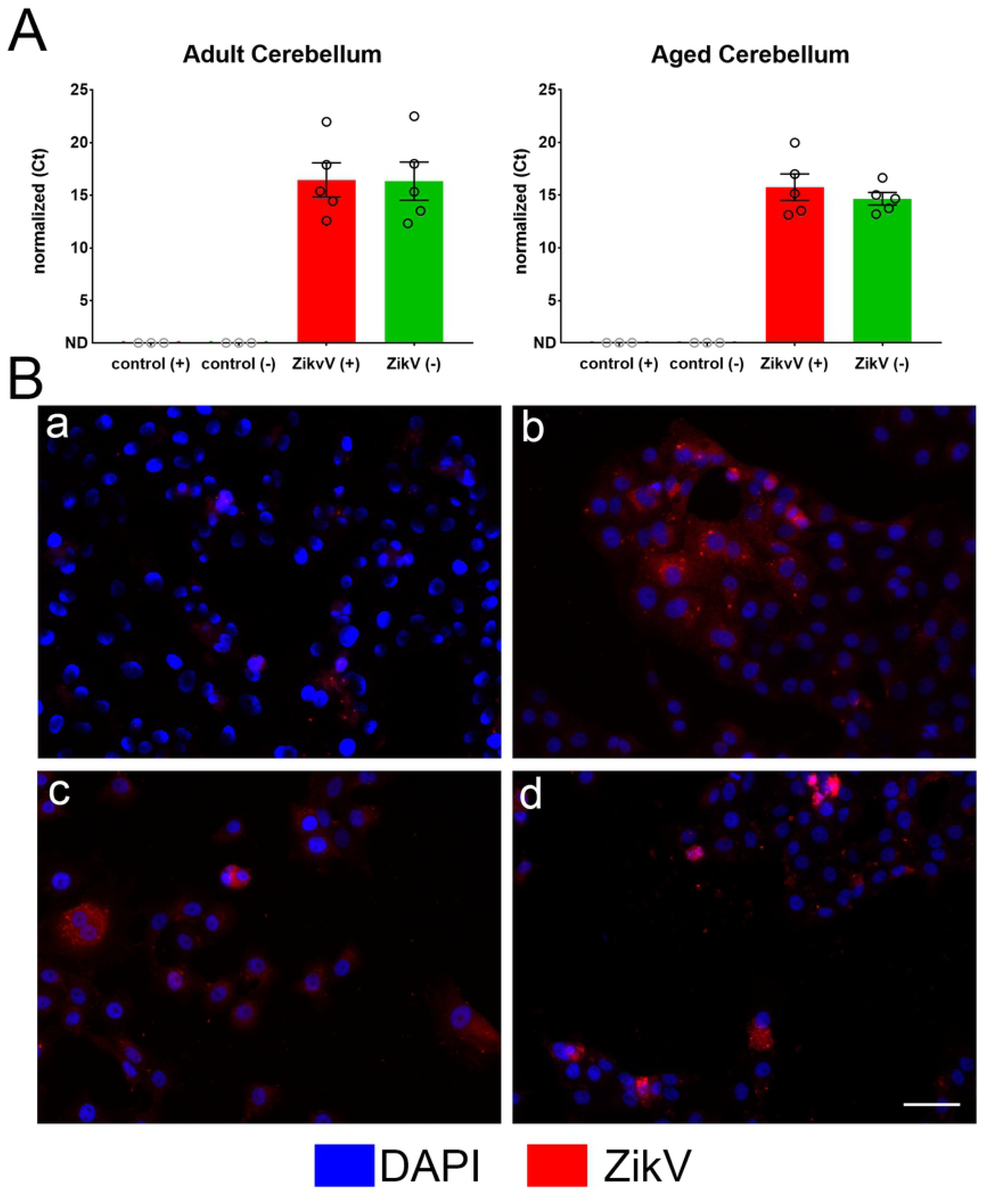
Evidence of live ZikV infection in adult and aged brains. **A**. ZikV-positive (red bar) and -negative sense RNA (green bar) as detected in RNA isolated from the cerebellum of uninfected (control) and ZikV-infected animals > 1 year old, indicating ZikV replicative intermediates in the brain. Ct values normalized to housekeeping gene (GAPDH). ND = no detectable amplification after 40 cycles. **B**. Infection of Vero E6 cells with cerebellar homogenate from (a) control mice (b) PRVABC059 (0.01 MOI as positive control). (c/d) cerebellar homogenate from 2 mice at 1ypi. All cultures were stained with anti-ZikV E-protein antibody and DAPI to mark cell nuclei. Scale Bar = 50μm.

## Discussion

The data presented above used immune competent mice exposed systemically to a clinically relevant dose of virus to explore the acute and long-term neurodevelopmental impact of Zika infection early in life. It shows that a peripheral (subcutaneous) challenge with ZikV shortly after birth causes a meningoencephalitis characterized by the infection of neurons in cerebellum, hippocampus, and the cerebral cortex that is associated with extensive apoptosis and immune cell infiltration, as well as reduced number of Purkinje cells and other GABAergic and granular neurons. The severe ataxia and dystonia that are evident during the acute infection subside within 30 days of challenge, and the animals appear to recover without overt motor or behavioral deficiencies. However closer examination, pairing MRI scans, behavioral testing, and gene expression studies shows that convalescent mice have reduced cerebellar volume and the magnitude of the defect correlates with deficits in motor coordination, hyperactivity, and impulsive or risky behavior that are evident throughout the life of the mouse. Further, we show that while the most severe tissue damage appears to occur during acute infection, the brain parenchyma in convalescent mice harbors low levels of replicating virus localized in small foci with active apoptotic processes and cellular infiltrates that are surrounded by activated astrocytes and microglia. The lesions are associated with a pattern of gene expression that indicates a continuous inflammatory milieu as late as a year post-infection. Together, these data suggest that subjects exposed to Zika early in development may not only experience long-term sequelae associated with the tissue damage that occurs during acute infection, but also harbor live replicating virus in their CNS that leads to continuous local inflammation and may worsen their long-term prognosis.

The most severe cases of CZS can include severe malformations of the brain (microcephaly, simplified gyral pattern, cerebellar hypoplasia, calcifications, ventriculomegaly), eyes (microphthalmia), and limbs and are diagnosed in 5-10% of children exposed *in utero* early in pregnancy. Further, there is mounting evidence that some infants exposed to the virus who do not exhibit CZS at birth, develop neurodevelopmental abnormalities such as hydrocephalus, or severe hearing and ocular defects later in life (27). Indeed, while initially ZikV was thought to be neurotropic only in early pregnancy, recent studies demonstrate astrogliosis, multifocal meningeal lesions, and altered neurodevelopmental processes in late pregnancy and even in infants, which is not surprising given that synapse formation, pruning and myelination processes extend for years after birth (28, 29). Further, recent *in vitro* studies show that while the virus preferentially infects immature progenitors, it can replicate in mature neurons (30, 31). Together, this raised concerns that, in addition to the expected sequelae of severe malformations due to early ZikV infection, perinatal exposure to ZikV, even if apparently asymptomatic, can cause subtle changes in neuronal maturation and neuroplasticity leading to motor, behavioral, learning, or psychiatric difficulties that may become evident years after exposure (27).

Developmentally, the CNS of mice at P1-3, like humans in week 23-32 of pregnancy, are undergoing extensive neurite outgrowth and synaptogenesis, have pre-oligodendrocyte progenitor cells that are actively undergoing proliferation, and a BBB that is established but not fully mature (32). ZikV challenge at this time resulted in infection of glutamic acid decarboxylase (GAD67+) neurons in the hippocampus, cerebral cortex, and most prominently the molecular layer of the cerebellar cortex(16, 20) with extensive neuronal loss. Correlation of the anatomical sequelae as detected by MRI, with histopathological changes and immunological responses over the lifespan of the animals shows that mice do not develop microcephaly or obvious calcifications or increased lateral ventriculomegaly. Increased fluid volume in the CNS, particularly in and around fourth ventricle and pia layers of the cerebellum and reduction in cerebellar size where most apoptotic lesions are also located during acute infection, are indicative of persistent changes to the microstructural composition of the tissue in this region. Importantly, the magnitude of the reduction in cerebellar volume correlated not only with the reduced latency to fall on the rotarod, but with the degree of hyperactivity and impulsivity. The results parallel a previous study conducted by Snyder-Kelley *et al* using young adult mice (60dpi) in a similar model and extend the observations to adult and aged mice demonstrating that although the virus retracts over the first 60 dpi, the neurological sequelae are permanent (33). Of note, increased fluid and/or disruptions in fluid diffusion in CNS have been described in patients with multiple neurologic and psychiatric disorders including children exposed in utero to HIV (34); future studies using higher resolution MRI in convalescent patients may provide some insight into ZikV-induced defects in infected children.

Immune responses in the CNS infection require a delicate balance between efficient pathogen clearance and the preservation of sensitive, non-renewable neural cells(35, 36). During acute ZikV infection, the majority of CD45+ infiltrating cells are T cells, particularly CD8+, which have been shown to play a key role in viral clearance, and mice with defective T cell responses succumb to infection(37), (16, 20). However, T cells can mediate tissue destruction and excessive inflammation(38). Importantly, previous studies have reported the long-term persistence of pathogen-specific resident memory CD8+ T cells in CNS that adopt a quiescent state (39, 40),(41). In this model however, the presence of apoptotic cells including apoptotic neurons, and a pattern of gene expression that suggest the presence of activated microglia/macrophage (*C3, Tnf, Ccl5, Cxcl10*), ongoing antigen presentation *(β2m, H2-Eb1, Cd-86*), and activated T cells (*Cd3ε, Cd8, Cd4, Ifn-γ, GrzmB, Prf1*, but not *Ctla4* or *Pd1*) as late as a year post infection suggests that the cells infiltrating these foci are not quiescent, but actively responding and possibly controlling the infection. Indeed, our preliminary studies suggest that the T cells that can be found in the CNS at 1ypi, adopt the phenotype of CD8+ resident effector memory cells. Future studies will be needed to fully characterize the T cells that persist in CNS as these could contribute to the pro-inflammatory milieu and worsen the neurological damage (42, 43). Indeed, while the lytic effect of the virus during acute infection is likely the predominant determinant of the life-long lesions and behavioral changes, the presence of proinflammatory cytokine and chemokines may impair healing and continue to skew the formation and pruning of synapsis contributing to the motor and behavioral defects observed(44). This could increase the risk of neurodegenerative conditions including Alzheimer’s, Parkinson’s, and multiple sclerosis over time(41, 45).

Many neurotropic viruses have been shown to persist in CNS, and in some cases can lead to recrudescence later in life(36). The nonlytic cellular mechanisms that govern viral latency and reactivation of DNA viruses, such as HSV and VZV, are reasonably well understood, however less is known about the persistence of RNA viruses in the CNS(22). Detectable levels of genomic RNA in the mouse brain parenchyma has been described for West Nile, Sindbis, Lymphocytic choriomeningitis, Borna disease virus, rabies virus, and influenza A virus suggesting that RNA viruses can establish long-term quiescent infection in CNS neurons but persistence of infectious flaviviruses in CNS has not been previously demonstrated (36),(46). Unlike those viruses, ZikV appears to remain replication-competent, albeit at levels far below those of the peak infection and confined to complex structures that are surrounded by activated microglia and astrocytes. Whether these cells contain the replicating virus impeding its spread or reflect a localized response to the low level of apoptosis induced by the virus remains unknown. The IHC imaging of the CNS foci in adult mice suggest that at least some of the neurons are infected by the virus, however the density of neurons within the foci appears to diminish over time. Thus, while long-lived CNS neurons may be an ideal harbor for long-term infection, it is likely that other cells may be involved as well.

A critical question stemming from the discovery that live virus persists in the CNS, is whether it contributes to the motor and behavioral defects. While our studies show that mice that survive an infection with ZikV early in life develop life-long defects in their motor skills and altered behavior, the deficiencies do not appear to worsen with age, suggesting that the tissue damage that occurs during the acute disease is the main determinant of the clinical phenotype, however the contribution of the pro-inflammatory milieu to the persistence of symptoms is still unknown. Given the potential plasticity of the brain early in life, future studies will need to explore whether eradicating the virus from the CNS and avoiding the protracted exposure to a proinflammatory milieu allows for a better reconstitution of the tissue. Conversely, as the foci of virus are surrounded by activated glia, it is possible that exposure to immunosuppressive or anti-inflammatory therapies such as TNF inhibitors could limit the tissue damage, but could also facilitate the spread of the virus (47, 48). or induce recrudescence of the virus in the CNS as has been observed with neurotropic coronavirus JHM (49) (50). Our preliminary studies treating chronically infected mice with cyclosporin or dexamethasone to suppress the immune response in ZikV did not result in increased mortality. Future studies will need to explore the impact of therapeutic immune modulators on viral load and sequelae.

In summary, these studies confirm that mild perinatal infections can results in long term neurological sequelae and concordant behavioral deficits that persist for the life of the host. Further, they provide a new understanding of the potential impact of ZIKV infections of the CNS by demonstrating that convalescent mice harbor multiple small foci with viral antigen, active apoptotic processes in neurons, and cellular infiltrates that appear to be walled-in by activated astrocytes and microglia as late as 1-year post-infection. The detection of negative strand viral RNA in addition to the genome, and recovery of infectious virus from the brain of convalescent mice demonstrates for the 1st time that ZIKV can establish life-long persistence in the CNS. The model described herein opens a novel opportunity to understand the role of the host immune response in active containment of the virus within the CNS and provides a platform to examine the role of the host immune response in active containment of the virus within the CNS and understand what circumstances could lead to viral recrudescence.

## Materials and methods

### Mice

C57BL/6 (B6) mice used in this study were bred and housed in sterile microisolator cages under 12-hour day/night cycle. They were given food and water ad libitum in the specific pathogen-free, AAALAC accredited animal facility of the U.S. Food and Drug Administration’s (USFDA) Division of Veterinary Medicine (Silver Spring, MD). This study was carried out in strict accordance with the recommendations in the NIH guide for the Humane care and Use of Laboratory Animals and with the approval of the White Oak Consolidated Animal Use and Care Committee at US-FDA.

### Zika Virus PRVABC59 (ZikV) Preparation

Zika virus PRVABC59 (Puerto Rico strain) used in this study is a contemporary strain that was isolated by the CDC from the serum of a ZIKV infected patient who travelled to Puerto Rico in 2015. The complete genome sequence is published (Ref. Gene bank accession # KU501215). Virus stocks from the CDC were kindly provided by Maria Rios (USFDA). This virus stock was expanded in Vero E6 cells. Briefly, cells were inoculated with the CDC provided stock for 1 hour at 37 °C, then provided growth media and incubated for 5-7 days at 37 °C, 5 % CO_2_. Supernatants were collected, clarified by centrifugation (10000 x g, 10 min) and the titer of the stock determined by TCID_50_ assay as previously described (51).

### ZIKV Infection and organ collection

All newborn mice were inoculated with 2000*TCID_50_ of ZikV, in 10 μl by subcutaneous (s.c.) injection with a 31G needle at the scruff of the neck. C57BL/6 mice (gender indeterminant) were inoculated one day after birth (P1). Control animals were injected (s.c.) with the same volume of diluent (MEM). All littermates received the same treatment in order to avoid cross-contamination. At the indicated time post-infection, mice were euthanized by CO_2_ asphyxiation and exsanguinated by transcardiac perfusion with ice-cold PBS prior to collecting the brains. For some studies, brains were bisected sagittally prior to freezing. Mice were weaned 21 days after birth and housed with same-sex littermates (2-5 mice per cage).

### Virus quantification by real-time qPCR

One microgram (μg) of isolated total RNA from infected and age-matched uninfected controls (procedure described below) was analyzed to establish ZIKV RNA levels, which are expressed as ZIKV copies/μg. Absolute quantification was determined using genomic viral RNA transcribed from a plasmid using the T7 Megascript transcription kit (ThermoFisher). The number of copies was calculated based on absorbance at 260 nm. One-step reverse transcription (RT)-real time qPCR was performed on a 10-fold serial dilution of this standard RNA in parallel with 1 μg of unknown sample RNA. This assay amplifies ZIKV genome position 1087 to 1163 based on ZIKV MR 766 strain (GenBank accession no. AY632535) Zika virus RNA transcript levels in the samples (52), in a single tube using a 25 μl reaction volume (RNA to cDNA kit, ThermoFisher). Reaction parameters for the Applied Biosystems Viia7 real time PCR machine: 60 °C for 30 min for reverse transcription, followed by 95 °C for 15 min to inactivate the RT enzyme and activate the DNA polymerase followed by 45 cycles of 95 °C, 15 sec and 60 °C for 1 min.

Detection of positive-and negative-sense strands of ZikV was performed as previous described (53) in a 2-step real-time PCR reaction. Briefly, reverse transcription of total RNA isolated from brain was performed using tagged ZikV-specific primers, combined with Oligo-dT primers for first-strand synthesis (High Capacity cDNA reverse transcription kit (ThermoFisher Scientific, Carlbad, CA). These reactions synthesized T7-tagged positive-sense and GVA-tagged negative-sense viral cDNA. The resulting cDNA was amplified using real-time qPCR with forward primers specific to the strand-specific Tags (T7 or GVA) and reserve primers specific to ZikV E gene sequences. A common Taqman probe specific for the ZikV amplicon was used to detect amplified cDNA. All custom primer/probes in this assay were produced using sequences previously described ((53), Table S2) and produced by Eurofins Genomics, USA.

### Taqman Low Density Array for gene expression

Flash-frozen brains (half brains bisected along the longitudinal fissure) were homogenized in 1 mL Trizol reagent (ThermoFisher, Carlsbad, CA) using a Precellys-24 (Bertin Instruments) equipped with a Cryolys cooling unit, with 1 mm Zirconia beads (BioSpec Products). RNA was isolated following the manufacturers’ protocol and resuspended in ultra-pure ddH_2_O. The concentration and purity of isolated RNA was determined by spectrophotometry at 260 nm and 280 nm using a NanoDrop spectrophotometer (ThermoFisher, Carlsbad, CA). To eliminate potential genomic DNA contamination, the DNA-free Turbo kit (ThermoFisher, Carlsbad, CA) was used as per the manufacturer’s protocol. Reverse transcription was performed on 1 μg of total RNA, using Multiscript High Capacity Reverse Transcriptase (ThermoFisher, Carlsbad, CA), per the manufacturer’s protocol, using random primers. The resulting cDNA was diluted ten-fold with ultra-pure, nuclease-free water and stored at −20 °C prior to use in real-time Taqman PCR reactions (ThermoFisher, Carlsbad, CA). The resulting cDNA was loaded into a Mouse Inflammation array card (ThermoFisher, Carlsbad, CA) and real-time qPCR was performed using a Viia7 real-time PCR instrument. All fold-change calculations were performed using the ΔΔCt method (54), normalizing against GAPDH and age-matched uninfected controls.

### Nanostring analysis of gene expression

Mouse Immunology Panel (v2) (Nanostring Technologies, Seattle, WA) Reporter and Capture probes were mixed with 100 ng of total RNA (20 ng/μl in molecular grade, ultra-pure ddH_2_O) isolated from the CNS of uninfected and ZikV infected animals. The hybridized probe/samples were then processed, and data acquired using the Nanostring MAX acquisition system, counting 280 fields of view (FOV) per sample. These data were normalized to internal controls and housekeeping genes in the panel. Gene expression is expressed as fold change in gene expression relative to age-matched, mock-infected control animals. The data was analyzed using Nanostring nSolver (ver. 4) and plotted using GraphPad Prism (ver 7).

### Immunofluorescence Immunohistochemistry (IF-IHC) and histology

Brain halves from ZikV-infected and age-matched uninfected control were placed in freshly prepared 4 % paraformaldehyde (PFA) solution (in 1 x PBS pH 7.4) immediately after extraction. After 24 hours of fixation at 4 °C, the tissue was transferred to 30 % sucrose in 1 x PBS for cryoprotection. The brains remained in 30 % sucrose at 4 °C until they sank in the solution (typically ≥ 24 hours). Sucrose embedded tissue was mounted in TissueTek O.C.T (Sakura-Finetek, Torrance, CA) and frozen. The brains were stored at −80 °C until sectioned. Sagittal sections (25 μm thick) were cut using a Leica CM1900 cryostat at a chamber temperature of −20 °C (Leica Biosystems, Buffalo Grove, IL) and mounted onto Superfrost-Plus Gold microscope slides (Fisher Scientific, Carlsbad, CA). Mounted sections were stored at −20 °C until staining. Prior to staining, sections were warmed to room temperature (RT) to dry, then immersed in phosphate buffered saline (PBS). For antigen retrieval, the sections were immersed in 0.1M Sodium Citrate solution, pH 8.5 at 80 °C for 8 minutes, then allowed to cool at room temperature for an additional 10 minutes. The sections were then washed with PBS and permeabilized using 0.5 % Triton X-100 in PBS for 60 minutes at RT. The sections were then blocked with 5 % normal goat serum and 1 % bovine serum albumin (BSA) in PBS + 0.5 % Triton X-100 for 2 hours. Primary antibodies used include: human polyclonal anti-Zika virus antibody (Kerafast EVU302, MA) or mouse anti-ZikV E protein mAb (Fitzgerald, 2713), neurofilament heavy chain cocktail (SMI31 and SMI32 mAb, Biolegend), rat anti-CD45 (BD Biosciences), rabbit anti-Ionized calcium-Binding Adapter molecule-1 (Iba-1) (Wako), rabbit anti-glial acidic fibrillary protein (GFAP)(Dako), mouse anti-glutamic acid decarboxylase 67 (GAD67) (SigmaMillipore). Tissue sections were incubated overnight in a humidified chamber at room temperature with primary antibodies diluted with 1 % BSA in PBS + 0.5 % Triton X-100. The slides were then washed with PBS + 0.05 % Triton X-100 followed by 1 x PBS and treated with the appropriate AlexaFluor (AF)-conjugated secondary antibodies (raised in goat) (ThermoFisher, Carlsbad, CA), diluted in 1 % BSA in PBS + 0.05 % Triton X-100 for > 120 min at RT. To increase sensitivity of the staining assay, we used the AlexaFluor 647 Tyramide Superboost (goat anti-rabbit) kit (ThermoFisher, Carlsbad, CA) as per the manufacturers’ protocol for detection of anti-ZikV E protein.

Staining for apoptosis was performed using the TUNEL assay (ApopTag-FITC, SigmaMillipore) preceded by antigen retrieval (as described above) as per manufacturer’s instructions. All IF-IHC sections were mounted with Pro-Long Diamond anti-fade mounting media containing DAPI (ThermoFisher, Carlsbad, CA). AF-labelled antibodies were detected at emission wavelengths: 405 nm (DAPI), 535 nm (Alexa-fluor 488), 605 nm (Alexa-fluor 568) and 650 nm (Alexa-fluor 647). Sections were imaged using an Olympus VS-120 virtual slide microscope (Olympus LSS) using a 20x or 40x objective lens. Regions of interest were captured using VS-Desktop software (ver 2.9, Olympus LSS). For confocal imaging, images were acquired using a Zeiss LSM 880 confocal microscope, using 405 nm (DAPI), 488 nm (AF488), 561 nm (AF568) and 633 nm (AF647) excitation lasers. Optimal fluorescence detection settings were determined and applied to all sections equally. Images were acquired using the Zeiss Zen software. Maximum intensity projections of acquired Z-stacks were compiled using Zeiss Zen and/or ImageJ + Fiji (NIH) software (55). Individual images were combined into figures using Adobe Photoshop CC.

### Behavioral Testing

Independent groups of animals began behavioral testing at 60 days post infection (young) or 90-180 days post infection (Adult) or 12-14 months post infection (aged adults). Mice were tested on the rotarod apparatus, followed by open field test (OFT), Novel Object Recognition (NOR), elevated plus maze test (EPM) and finally the social interaction (SI) test, in this order. Each group underwent 1 test per day. For adult and aged groups, there was a six-week delay between NOR and EPM tests due to equipment availability. All mice were tested blindly so the tester did not know the age or prior infection status of the mice.

#### Rotarod

Locomotor coordination and endurance was assessed using the rotarod test (IITC Life Science; Woodland Hills, CA). Mice underwent 3 consecutive 180 s trials, during which they were placed on a rotating drum (1 1/4 in diameter) that accelerated from 5 rpm to 20 rpm. At least 2 independent reviewers monitored the session and recorded the time the mouse fell off the drum. If the mouse did not fall, a time of 180 s was recorded. The average latency time to fall for the three trials was used for analysis.

#### Open Field Test (OFT)

Horizontal exploratory locomotor activity and anxiety behavior were assessed using the OFT. Mice were placed in the corner of a square arena (40 cm x 40 cm) and allowed to explore freely for 30 min. The trial was recorded with an overhead camera; total distance traveled and time spent in the center of the arena (interior 50%) was automatically scored using TopScan software (CleverSys; Reston, VA) or AnyMaze Software (Stoelting Co., Wood Dale, IL)

#### Elevated Plus Maze (EPM)

Anxiety-like behavior was assessed in the EPM test. The EPM (Stoelting, Inc; Wood Dale, IL) is a raised maze (50 cm above the ground) with two enclosed arms (35 × 5 x 15 cm) perpendicular to two open arms (39.5 × 5 cm) intersected by an open central area (5 × 5 cm). Mice were placed in the center of the maze facing one of the two open arms and recorded with an overhead camera as they freely explored the maze for 10 min. Mice that fell off the maze were immediately returned to the same arm in the same position. Time spent in the open arms and total distance traveled was determined using TopScan software (CleverSys; Reston, VA) or AnyMaze Software (Stoelting Co., Wood Dale, IL)

#### Novel Object Recognition (NOR)

For this test, animals are pre-conditioned one day before the test, by placing the animals in the open field test box, with 2 identical objects (Black spheres or white cubes). The mice freely explore these items for 10 minutes and were then returned to their home cages. To test for novel object recognition, on the next day one of the 2 objects was exchanged for a novel object (eg/ if pre-conditioned with 2 white cubes, then exposed to 1 white cube and 1 black sphere). The animals were then returned to the open field test box for 5 minutes. The trial was recorded with an overhead camera. Time spent with the familiar and novel objects, as well as distance travelled, were automatically scored using TopScan software (CleverSys; Reston, VA) or AnyMaze Software (Stoelting Co., Wood Dale, IL)

#### Social Interaction (SI)

Sociability and social novelty preference were assessed in the SI test as described (Clark et al, 2018). The SI arena (Stoelting, Inc) is a 40 × 40 cm chamber divided into 2 equal sized arenas (18 × 20 cm), connected at their base by a smaller arena (11 × 20 cm) used to traverse between them. Two empty small wire holding cages, weighted at the top, were placed in the far corners of the 2 large arenas. After experimental mice were habituated to the chamber (5 min), previously habituated conspecific, stranger mice were placed in one of the 2 holding cages. Experimental mice then freely explored for 5 min while being recorded with an overhead camera. Total locomotion and time spent near (within 4 cm) the conspecific mouse was determined using TopScan software.

#### Magnetic Resonance Imaging (MRI)

All MRI experimental measurements were performed with an Agilent 7T (300 MHz) horizontal bore small animal MRI (Agilent Inc.; Santa Clara, California) with an open bore of 310 mm and a diameter of 115 mm inside the gradient coil (Resonance Research Inc.; Billerica, MA). The specification of the transmit birdcage RF coil (RAPID MR International Inc.; Columbus, Ohio) is; quadrature driving 12-rung birdcage coil without an RF shield, inner diameter (ID) = 72 mm, inner length = 110 mm, outer length = 120 mm, end-ring width = 5 mm, and rung width = 5 mm. All animals were maintained on 2-3 % isoflurane during the tests, while vital signs were being monitored to ensure minimal pain and distress. A spin echo multi-echo multi-slices imaging sequence was used for the in-vivo experiment with TR/TE = 850/10 msec, Number of echo = 5, Number of average = 3, Dummy scan = 0, Data matrix (RO x PE) = 256×256, FOV = 50 × 50 (for Sagittal image) and 40 × 40 (for Transverse image) mm^2^, Number of slices = 15, Thickness = 1 mm, Gap = 0 mm, and Scan time = 10 min. 53 sec. The 3D volume mapping and quantitative analysis of fluid volume from acquired MR images were conducted using an open source software, 3D Slicer (ver 4.10)(www.slicer.org; (56)).

### Statistical Analysis

Statistical analysis for behavioral studies were conducted using both R (R Core Team (2019). R: A language and environment for statistical computing. R Foundation for Statistical Computing, Vienna, Austria. https://www.R-project.org/.) and GraphPad Prism (v. 8.0.1: San Diego, CA). T tests, ANOVA or Mann-Whitney non-parametric tests were used to determine the overall effects of Zika on cytokine levels, behavioral changes and virus titers were used as appropriate. Mann-Whitney non-parametric tests were used to test the effect of Zika within age group. False discovery rate was used to identify gene expression that were significantly modified by the infection. Finally, Spearman’s rank correlation coefficient were used to correlate MRI changes, behavior testing and cytokine levels as appropriate. Data are shown as mean +/-SEM; *p* < 0.05 was considered significant. *p<0.05, **p<0.01, ***p<0.001.

## Acknowledgments

The assertions herein are the private ones from the authors and are not to be construed as official or as reflecting the views of the Food and Drug Administration. The authors wish to thank John Dennis, Damaris Molano, Mary Belcher, Claudia Diaz, Tina Smith-Curry and the personnel of the FDA Department of Veterinary Medicine. We thank Dr Bu Park for performing the MRI studies and analysis. We thank Christian Sauder, Dino Feigelstock, Ashutosh Rao, and Amy Rosenberg for reviewing the manuscript. We thank Dr Maria Rios for generously providing PRVABC59 (ZikV). This work was supported in part by a Center of Excellence in Regulatory Science and Innovation (CERSI) grant to University of Maryland from the US Food & Drug Administration (Grant: U01FD005946). This study was supported in part by Senior Postgraduate Research Fellowship Awards to A.L., H-N.L., I.M.W., K.E., and JS. from the Oak Ridge Institute for Science and Education through an interagency agreement between the U.S. Department of Energy and the U.S. Food and Drug Administration. Lastly, this work was partly supported by grants funded by the FDA’s Office of Counter-terrorism and Emerging Threats.

## Supplementary Data Legends

**S. Table 1: Correlations of MRI measurements and Behavior**. Spearman’s rank correlation were used to compare brain characteristics as measure during MRI volumetric analysis and the indicated behavioral parameters. Statistically significant correlations are indicated with blue, underlined values.

**S. Table 2: Correlations of inflammatory gene expression and Behavior**. Spearman’s rank correlations were used to compare immune gene expression and the indicated behavioral parameters. Statistically significant correlations are indicated with blue, underlined values.

**S. Figure 1: Cellular infiltration and inflammation in the CNS of convalescent ZikV-infected mice**. Sagittal sections from control and ZikV-infected brains at 12 and 30 dpi. **A**. Sections stained with Iba-1 (red) to mark microglia. Note activation of microglia through the brain at 12 dpi is significantly reduced by 30 dpi in the cerebral cortex, hippocampus and thalamic regions. More focal microglia activation is found in the cerebellum at 30 dpi. **B**. Sections stained with CD45 (green) to mark infiltrating immune cells. Images representative of 3-5 mice.

**S. Figure 2. Neuropathology in convalescent ZikV-infected mice. A**. Cerebellum of control and ZIKV infected mice stained with DAPI (Blue) and Neurofilament staining (green). Arrow indicates Purkinje cell layer. **B**. Regions in the cerebral cortex and hippocampus stain positively for apoptosis (TUNEL, green) and the activation of microglia (Iba-1+, red) at 22 dpi. These lesions are resolved by 30 dpi. dg = dentate gyrus, hc = hippocampus, OCx =Occipital cortex. Scale bars (A&B)= 500 μm.

**S. Figure 3. Behavioral data: A**. Weight of the animals tested in behavioural studies. **B**. Representative examples of heat tracking maps for 60 dpi mice recorded in the open field and Elevated Plus Maze tests (representative of 6-7 mice/group). The open arm of the maze is labelled. The opposite arm is closed. **C**. Novel Object recognition (NOR). Bars show interaction ratio (Novel:Familiar) as the time spent with new vs familiar objects **D**. Social Interaction Test. Data indicates total movement in the maze and Sociability index (SI, Ratio of Social:Non-social with mice in the maze). Statistical analysis between infected and uninfected groups: T test. * P<0.05

**S. Figure 4. ZikV infects neurons, including GABAergic neurons in the cerebellum**. DAPI (blue) stains nuclei in all images. Left column (green), cell specific staining for astrocytes (GFAP), neurons (Neurofilament, NF-H) and GABAergic neurons (GAD-67). Center column, staining for ZikV (red). Left column (Merge). White arrows indicate colocalization of virus with NF-H and GAD67. Representative image from 3-5 mice. Scale bars = 50 μm.

**S. Figure 5: Apoptosis and cellular infiltration in sagittal sections of brain:** Sagittal sections stained for apoptotic (TUNEL+, green), and immune infiltrating (CD45+, red) cells. Evidence of CD45+ cells and apoptosis in hippocampus and cerebellum convalescent animals at 60 dpi. Image representative of 6 animals.

**S. Figure 6. Apoptotic foci in cerebellum of ZikV convalescent mice. A**. Confocal image of ZikV+ foci in cerebellum of 1 ypi ZikV-infected mice, shows position of TUNEL^+^ cells (apoptosis, green) relative to Iba^+^ microglia (red) and infiltrating CD45^+^ cells (red). Note that microglia proximal to apoptotic lesions shows hypertrophy of the nuclei and short processes consistent with activation (arrow head), while those distant to the lesion show small nuclei and fine processes (arrow). CD45+ infiltrating lymphocytes are also proximal to apoptotic foci. **B**. Individual channels for: Apoptosis (Apop, green), astrocytes (GFAP, red), microglia (Iba-1, red) and infiltrating lymphocytes (CD45+, red) and merged images with orthogonal views, for apoptotic lesions also shown in Figure 5.

**S. Figure 7. ZikV RNA levels in the cerebral cortex of convalescent mice**. Single reaction taqman RT-qPCR for viral RNA performed on total RNA isolated from the cerebral cortex of ZikV-infected or age-matched control animals. A sub-population of animals demonstrated low levels of viral RNA in the cerebral cortex.

